# Enhancing Breast Ultrasound Segmentation through Fine-tuning and Optimization Techniques: Sharp Attention UNet

**DOI:** 10.1101/2023.07.14.549040

**Authors:** Donya Khaledyan, Thomas J. Marini, Avice O’Connell, Kevin Parker

**Affiliations:** Department of Electrical and Electronics Engineering University of Rochester, Rochester, NY, USA; Department of Imaging Sciences University of Rochester Medical Center, Rochester, NY, USA

**Keywords:** UNet, optimization, segmentation, breast ultrasound imaging, fine-tuning

## Abstract

Segmentation of breast ultrasound images is a crucial and challenging task in computer-aided diagnosis systems. Accurately segmenting masses in benign and malignant cases and identifying regions with no mass is a primary objective in breast ultrasound image segmentation. Deep learning (DL) has emerged as a powerful tool in medical image segmentation, revolutionizing how medical professionals analyze and interpret complex imaging data. The UNet architecture is a highly regarded and widely used DL model in medical image segmentation. Its distinctive architectural design and exceptional performance have made it a popular choice among researchers in the medical image segmentation field. With the increase in data and model complexity, optimization and fine-tuning models play a vital and more challenging role than before. This paper presents a comparative study evaluating the effect of image preprocessing and different optimization techniques and the importance of fine-tuning different UNet segmentation models for breast ultrasound images. Optimization and fine-tuning techniques have been applied to enhance the performance of UNet, Sharp UNet, and Attention UNet. Building upon this progress, we designed a novel approach by combining Sharp UNet and Attention UNet, known as Sharp Attention UNet. Our analysis yielded the following quantitative evaluation metrics for the Sharp Attention UNet: the dice coefficient, specificity, sensitivity, and F1 score obtained values of 0.9283, 0.9936, 0.9426, and 0.9412, respectively. In addition, McNemar’s statistical test was applied to assess significant differences between the approaches. Across a number of measures, our proposed model outperforms the earlier designed models and points towards improved breast lesion segmentation algorithms.

## 1. Introduction

Breast cancer is a major public health concern, ranking second in cancer-related deaths among women globally, and it is estimated that 1 in 8 females in the United States will develop breast cancer in their lifetime [1–4]. Early detection and treatment are vital for preventing metastasis and improving survival rates. Screening mammography has been successful in reducing mortality by allowing early detection [5]. However, most of the world currently lacks access to any form of medical imaging for the evaluation of breast masses [6–9]. Ultrasound is a low-cost and portable imaging modality that is first-line evaluation for palpable breast lumps, but its deployment has been limited by the ability to obtain ultrasound imaging which typically requires an expert sonographer to acquire the images and a specialist to interpret them. Artificial intelligence is a promising avenue to circumvent the issue of a lack of experienced sonographers. The value of breast ultrasound segmentation of lesions plays a potentially vital role in artificial intelligence, which could then be used to achieve these goals [10]. Accurately outlining and identifying regions of interest in ultrasound images could allow the development of practical artificial intelligence solutions which could aid in early detection and treatment [11, 12]. Additionally, artificial intelligence can be a replacement solution in situations and areas in which there is a lack of specialists; it can also aid specialists in better diagnosis.

Over the past two decades, extensive research has been conducted to develop effective approaches for achieving precise outlines of regions of interest. These approaches include various methodologies and techniques that are specifically designed to address the challenges inherent in breast ultrasound image segmentation [13–15]. Deep learning (DL) models, predominantly convolutional neural networks (CNNs), have proven remarkable success in accurately segmenting anatomical structures and identifying pathological regions in various medical imaging modalities, including X-ray, MRI, CT, and ultrasound [16–21].

With ultrasound’s safety, portability, cost-effectiveness, and non-invasiveness, coupled with robust machine learning (ML) algorithms, we can potentially enhance early detection, assessment accuracy, and treatment outcomes for breast cancer patients with the development of innovative artificial intelligence solutions. However, ultrasound images can include high variations in terms of image quality and artifacts [22]. Optimizing and fine-tuning the model enables it to adapt and perform well across different variations and enhances its applicability in real-world scenarios. Moreover, with the increase in data and model complexity, optimization and fine-tuning play a vital and more challenging role than before[23–26].

The fundamental role of an ML model is to configure the model or determine the algorithm used for minimizing the loss function. The choice of hyper-parameter values directly impacts the performance of the ML model[23].To achieve an optimal ML model, it is essential to explore a range of possibilities. This process, known as hyperparameter tuning [27], involves designing the ideal model architecture while finding the optimal configuration for the hyperparameters [23, 28].

In this study, we explore the impact of different optimization and fine-tuning techniques on the segmentation results of UNet based models; these techniques include but are not limited to the activation function, loss function, input size, batch size, weight initialization, learning rate schedules, and early stopping on the overall performance of the segmentation models. These techniques aim to improve convergence speed, alleviate overfitting and underfitting, and enhance the model’s generalization capabilities [24]. We also tested the performance of UNet, Sharp UNet, Attention UNet, and a novel combined approach called Sharp Attention UNet. We hypothesized that Sharp Attenuation UNet would have the best performance compared to the mentioned networks, because the combination of salient features as the outcome of attention gates and the sharpened features will improve the network’s ability to extract clinically important features.

The findings of this study are expected to provide valuable insights into the optimization process for breast ultrasound image segmentation models. By identifying the most effective combinations of activation functions, loss functions, and other optimization techniques the reliability of segmentation models will improve leading to better clinical decision-making and patient care.

## 2. Dataset and data preprocessing

### 2.1 Data collection

The dataset we used in this study is known as “Breast Ultrasound Images” (BUSI) [29]. The dataset consists of 780 images from 600 female patients between the ages of 25 and 75, collected in 2018. They were scanned using a LOGIQ E9 ultrasound system and lesions were segmented with manually outlined masks from the radiologist’s evaluation. The images are classified into three groups: (1) 133 normal images without masses, (2) 437 images with benign masses, and (3) 210 images with malignant masses. The images are in PNG format, have varying heights and widths, and an average size of 600*500 pixels. The data was preprocessed by removing non-image text and labels.

### 2.2 Image enhancement

The accuracy of DL models can be enhanced by employing image enhancement techniques prior to feeding them to the network [30, 31]. Ultrasound images often contain multiple artifacts such as speckle noise, attenuation effect, and low contrast, leading to diminished image quality and interpretation challenges. To address these issues, image enhancement techniques can be employed to mitigate or minimize these artifacts and enhance the overall image quality. Enhancing the image quality clarifies patterns which DL models are learned based on them, enabling the model to identify and classify features within the image more accurately.

In this study, we utilized contrast limited adaptive histogram equalization (CLAHE) [32] as an image enhancement technique. CLAHE is an improvement over adaptive histogram equalization (AHE) [33], and it mitigates the problem of excessive contrast levels in AHE. Unlike AHE, which can exceedingly enhance contrast, CLAHE sets a constraint on the contrast using a histogram. The objective of CLAHE is to enhance image contrast while preserving image quality. This process requires operation on localized image regions, referred to as “tiles” to align the contrast within each tile with a specified histogram shape. To achieve a unified and continuous output image, neighboring tiles are merged using bilinear interpolation. The CLAHE method can be formulated as eq. 1, where *β* is clip limit, *M* is area size, *N* is number of grey-level values, α is clip factor, and *S*_max_is maximum tolerable slope.

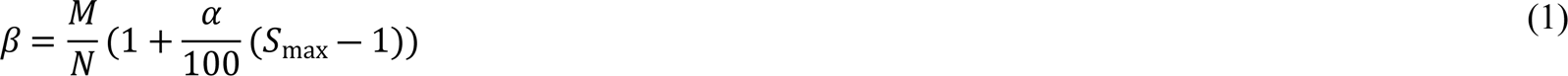

### 2.3 Data augmentation

Data augmentation is a technique that involves modifying the existing training data to generate new data samples, which helps to increase the size of the training dataset. This can reduce the likelihood of overfitting by introducing more variability and diversity in the training data results in improving the model’s performance and generality. Data augmentation aims to create a more diversified set of training data that better represents real-world scenarios and can help the model generalize better on new, unseen data, especially in the medical imaging field, where lack of data is a concern.

Though applying augmentation can help to diverse, generalize, and balance the data, it is crucial to consider the potential negative impact of the selected augmentation parameters on the diagnostic ability of the system. For instance, in ultrasound images, high brightness or zoom values may result in loss of important details or distortions that could affect the accuracy of the model predictions, as ultrasound artifacts such as shadowing or enhancing contain important information to diagnose the abnormality [34]. Therefore, it is essential to carefully select the appropriate augmentation parameters and adjust them to avoid any potential degradation of the image quality or loss of essential information.

Furthermore, it is worth noting that the specific choice of augmentation parameters can vary depending on the type of image data being used [35, 36]. For ultrasound images, it may be beneficial to use augmentation techniques that preserve the underlying structures and features of the images while introducing sufficient variability to prevent overfitting. For example, subtle rotations or shifts may be more appropriate than drastic changes in brightness or zoom level.

The parameters used in this study include 45-degree rotation, which rotates the image randomly in the range of [-45,45] degrees. The [-0.08,0.08] zoom range randomly zooms in or out of the image. The horizontal flip is on, which flips the randomly selected images horizontally. Moreover, width shift range and height shift range are both 0.15, it shifts the image horizontally and vertically, respectively. The shear range is [-0.03,0.03], it will apply the shear transformation to the image, and the brightness range changes between 0.99 and 1.07, which adjusts the brightness of the image. These parameters are used to define the data generators, which are functions that generate batches of augmented data samples during the training of an ML model. By randomly applying these parameter values to the training data, the resulting augmented dataset better represents real-world scenarios and reduces overfitting of the model to specific instances in the training set.

In the results section, we will present the segmentation model’s performance both with and without the utilization of the data augmentation technique.

## 3. Materials and Methods

### 3.1 Model Optimization Techniques

In this section, we focus on exploring various optimization techniques, including activation functions, dropout, and loss functions to fine-tune our segmentation model. By carefully selecting and fine-tuning these optimization components, we aim to improve the model’s ability to accurately outline masses in breast ultrasound images while identifying images with no mass.

#### 3.1.1 Activation function

Activation functions are essential parts of neural networks. Nonlinear activation functions allow neural networks to model complex nonlinear relationships between inputs and outputs, which is necessary for many real-world applications. They will ensure that gradients can be propagated through the neural network during backpropagation. The derivative of the activation function regarding its input defines the gradient flow through the network, and different activation functions can affect the stability and speed of gradient propagation.

Rectified Linear Unit (ReLU) is a simple and widely used activation function that returns the input value if it is positive, and zero otherwise [37]. It has been shown to be effective in many neural network architectures due to its simplicity and speed. However, one potential downside of ReLU is that it can suffer from the “dying neurons” problem, where a large portion of the network’s neurons can become non-responsive and “die” during training. To address the “dying neurons” problem, the Leaky ReLU (LReLU) activation function was introduced. This function is similar to ReLU but returns a small negative value instead of zero for negative inputs. This ensures that all neurons are active during training, which can lead to better performance. However, with all the advantages over ReLU and the previous activation functions, LReLU still has a sharp edge at 0 value. This can cause optimization issues during training, particularly for gradient-based optimization methods. The sharp edge can lead to abrupt changes in the function’s output, which can result in vanishing or exploding gradients. These problems can make the optimization process unstable, slow down training, or even prevent the model from converging altogether.

To overcome this problem, swish was proposed. Swish is a newer activation function that has gained popularity in recent years. It was invented at Google Brain in 2017 by Ramachandran et al. [38]. It is a smooth, non-monotonic function that is similar to the sigmoid function. Swish has been shown to outperform ReLU and LReLU in some neural network architectures, but its performance can be sensitive to hyperparameters. While swish has shown effective performance on certain datasets and architectures, it is not a universal solution that works well for all neural networks and tasks. Swish activation function’s equation is:

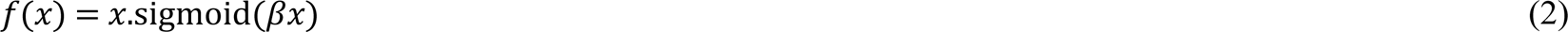

which *β* is a learnable parameter. However, most implementations exclude the use of this trainable parameter, resulting in the activation function being:

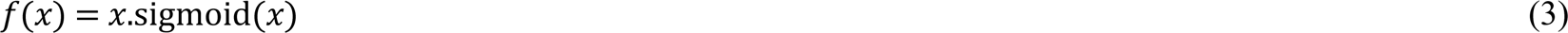

where sigmoid(*x*) is:

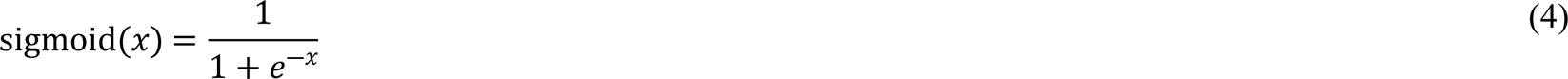

Another novel activation function that has demonstrated effectiveness in various applications is mish. Similar to swish, mish is a non-monotonic activation function for deep neural networks, proposed by Diganta Misra in 2019[39].

The mish activation function is defined as eq.5:

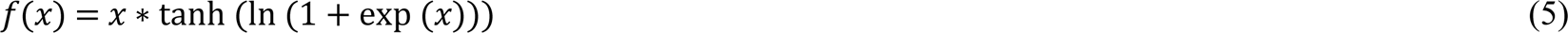

where *x* is the input to the activation function. The mish function is a smooth, continuous, and non-monotonic function that is symmetric around the origin. It has a maximum value of 1.0 at *x* = 0 and asymptotes to linearity for very small and very large values of *x*.

The mish activation function offers several advantages when compared to other activation functions. One advantage is its ability to enhance the performance of deep neural networks while exhibiting self-regularization. This property mitigates the risk of overfitting by managing the growth of gradients during training. Additionally, the mish activation function is computationally efficient and straightforward to implement, requiring only a small number of elementary operations.

As illustrated in Figure 1 (a) mish and swish both possess non-monotonicity, smoothness, and the ability to retain a small quantity of negative weights. These characteristics are responsible for the reliable performance and enhancement of deep neural networks. In the optimization process of DL networks, the first and second derivatives of the activation function can provide critical information about the shape and direction of the function. By analyzing these derivatives, we can determine the direction of steepest descent and whether the function is concave or convex. In the case of mish and swish activation functions, their distinctive negative curvature and smoothness, as evidenced by their first and second derivatives in Figure 1 (b), allows for more efficient optimization during the gradient descent process. This can lead to faster convergence and better performance of DL networks.

**Figure 1.**
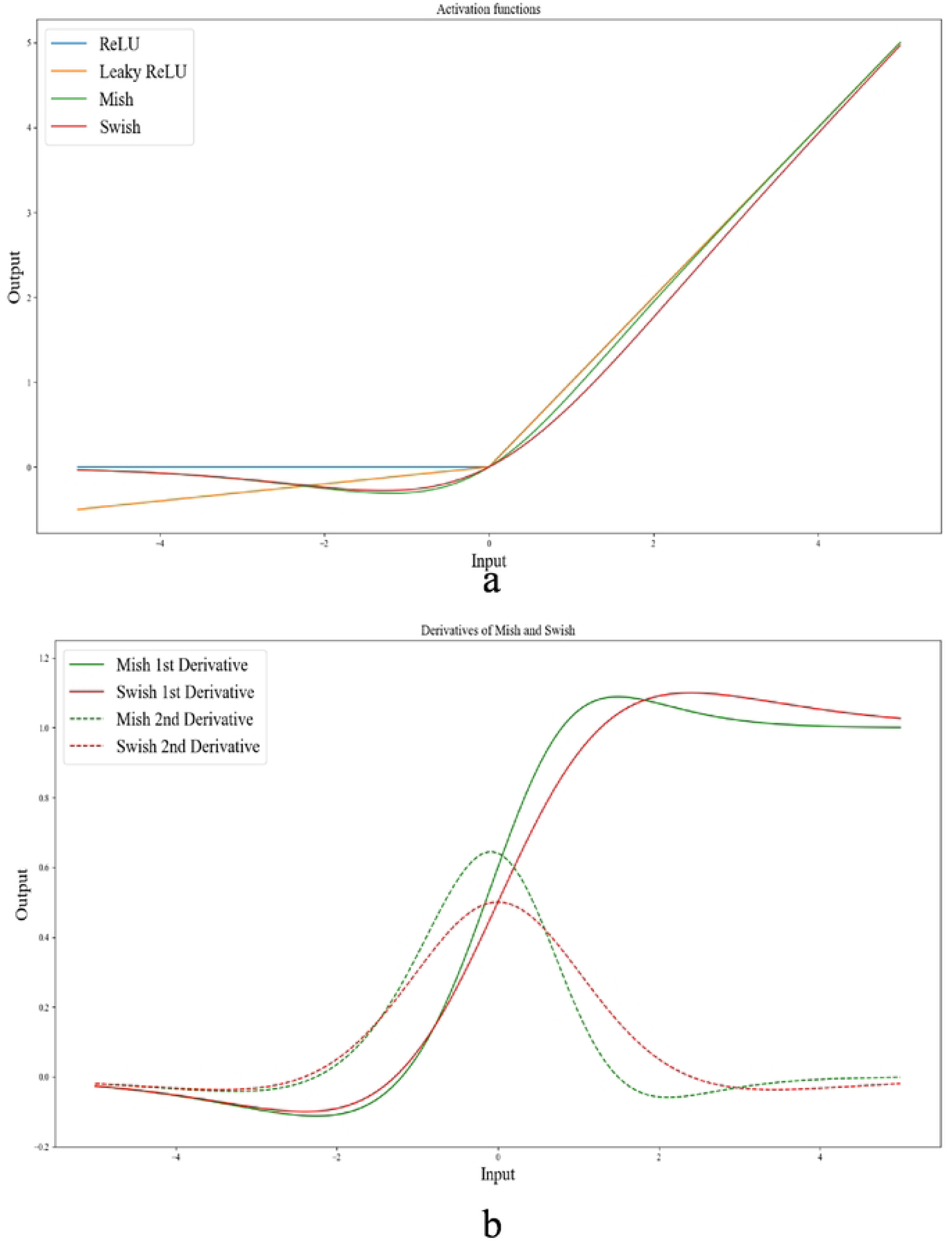
(a) Graph of ReLU, Leaky ReLU, mish, and swish. Both mish and swish have a unique negative curvature, setting them apart from ReLU, and Leaky ReLU. b) The 1^st^ and 2^nd^ derivatives of mish and swish activation functions (graphs are plotted in VS Code).

#### 3.1.2 Dropout

Dropout is a regularization method exploited in neural networks to reduce overfitting. Overfitting occurs when a model becomes too complicated compared to the number of data available for training and starts to fit noise in the training data, rather than the underlying patterns [40]. This leads to poor generalization and high error rates on new, unseen data [41].

Dropout avoids overfitting by randomly dropping out a fraction of the neurons in a layer during training. This drives the remaining neurons to learn more robust features that are not dependent on the presence of specific neurons. By doing so, the model becomes less sensitive to the specific details of the training data and is more likely to generalize well to new data. Furthermore, dropout can reduce the effect of co-adaptation between neurons. When neurons co-adapt, they tend to learn analogous features and can become excessively specialized, which can lead to overfitting. There have been many studies regarding the optimal values for dropout. Selecting the optimized drop out values depends on various factors such as the dataset, network architecture, and training method used.

Gal et al. [42] explored the impact of dropout probability and the number of neurons on the performance of deep neural networks. They suggested that higher dropout probabilities are generally better for deeper layers of the network and that the optimal dropout probability may vary based on the specific task and dataset. The optimal value for dropout in this work is 0.1 for the encoder and 0.5 for the decoder. These dropout values were selected based on the recommended optimal values provided in reference [42] and were further fine-tuned through an iterative process of trial-and-error around these values.

#### 3.1.3 Loss function

The loss function utilized in this study is a custom loss function for training neural networks for segmentation tasks (eq.6). It combines two loss functions, the Binary Cross-Entropy (BCE) loss and the dice loss. The BCE loss is a common loss function deployed for binary classification problems, such as image segmentation, where each pixel is classified as either foreground or background. The *BCE* loss measures the difference between the predicted probabilities and the true binary labels (eq.7).

The Dice loss, on the other hand, measures the overlap between the predicted segmentation mask and the true mask. It is calculated as eq.8, where the Dice coefficient (eq.9) measures the similarity between the predicted and true masks[43].

By combining these two loss functions, the model is encouraged to produce segmentation masks that are both accurate and have high overlap with the true masks. This can lead to better performance on segmentation tasks, particularly when dealing with complex or ambiguous boundaries between foreground and background classes.

The equations for *BCE*_Dice_loss is defined as follows:

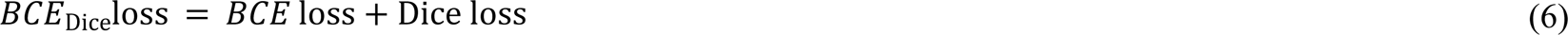

where *BCE* loss is calculated as:

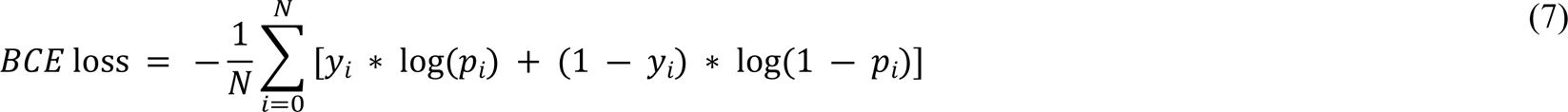

where *y* is the ground truth label (either 0 or 1), *p* is the predicted probability of the positive class (i.e., the probability of the pixel being part of the object), and log is a natural logarithm function.

The Dice loss is calculated as:

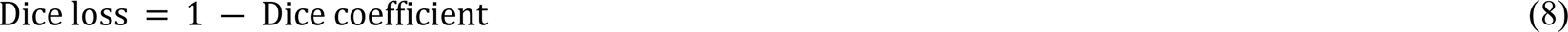

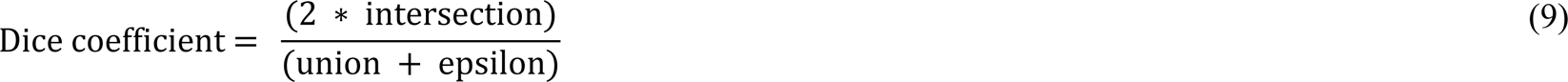

where intersection is the number of pixels where the predicted and ground truth masks both have a value of 1, union is the number of pixels where either the predicted or ground truth mask has a value of 1, and epsilon is a small value (e.g., 1e-5) to avoid division by zero.

From the equations, with *BCE* loss we can handle training when the foreground and background classes are imbalance, and since the size of the segmented area of the lesions is smaller than background, *BCE* loss is an appropriate way to tackle that. In the other hand, Dice loss is more sensitive to the foreground which means segmentation accuracy, that would be rational to take advantage of that for an appropriate segmentation. Thus, *BCE*_Dice_loss was employed in this study because it combines these two terms to penalize false positives and false negatives (through the *BCE* loss term) while also encouraging overlap between the predicted and ground truth masks (through the Dice loss term). The final *BCE*_Dice_loss is the sum of these two terms.

### 3.2 Models and Network architectures

In this section, we explore the models we used with the optimization and fine-tuning steps introduced earlier. Each architecture in Figure 2, 3, and 4 are designed based on the highest score hyperparameters and network fine-tunings. The backbone of the architectures explored in this section are based on UNet.

**Figure 2.**
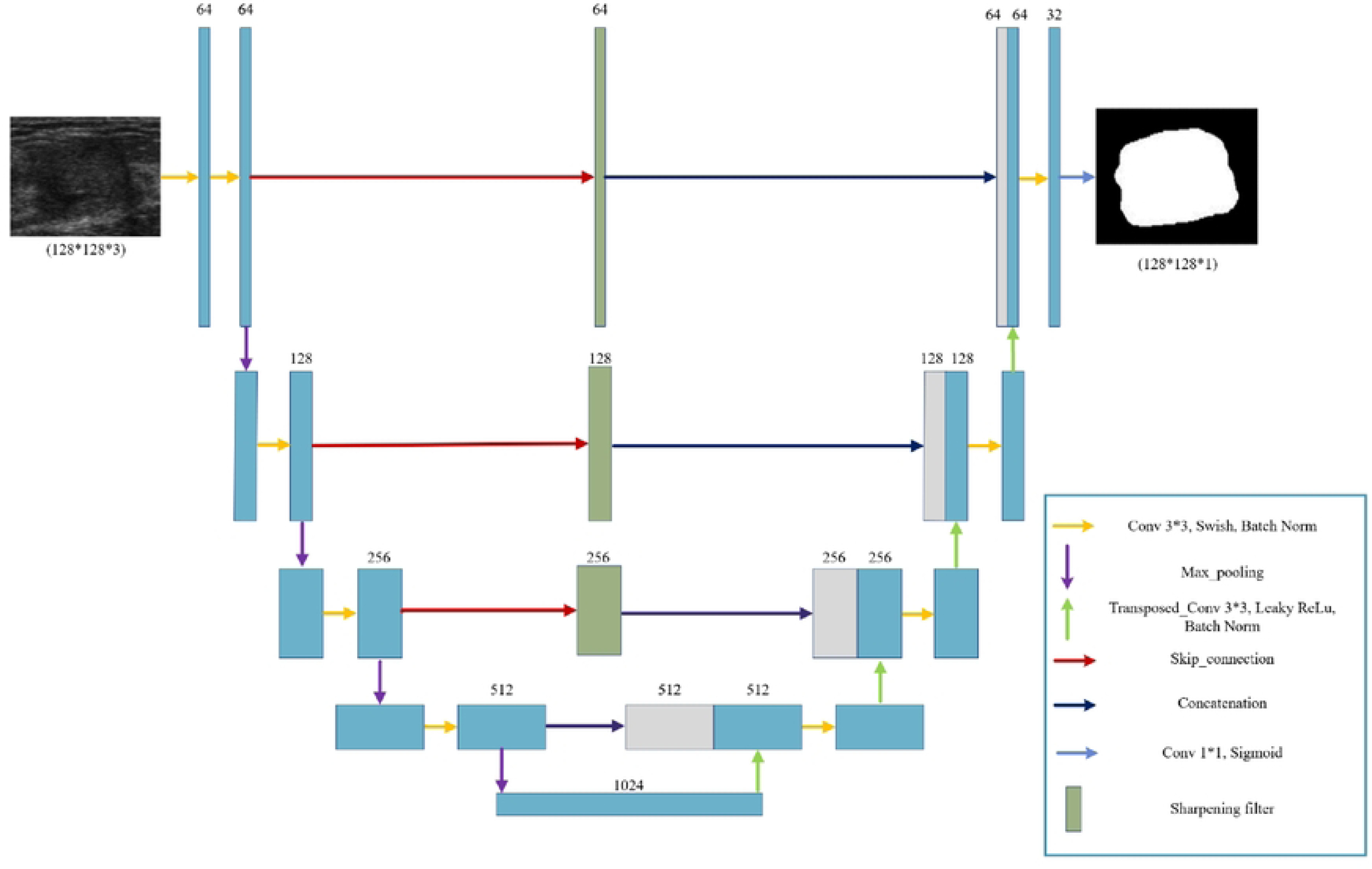
The Sharp UNet architecture inspired by [48] includes a schematic layout where encoder features are convolved with a sharpening spatial kernel before merging with the decoder features. This helps to reduce feature mismatches without adding extra parameters or computational cost.

**Figure 3.**
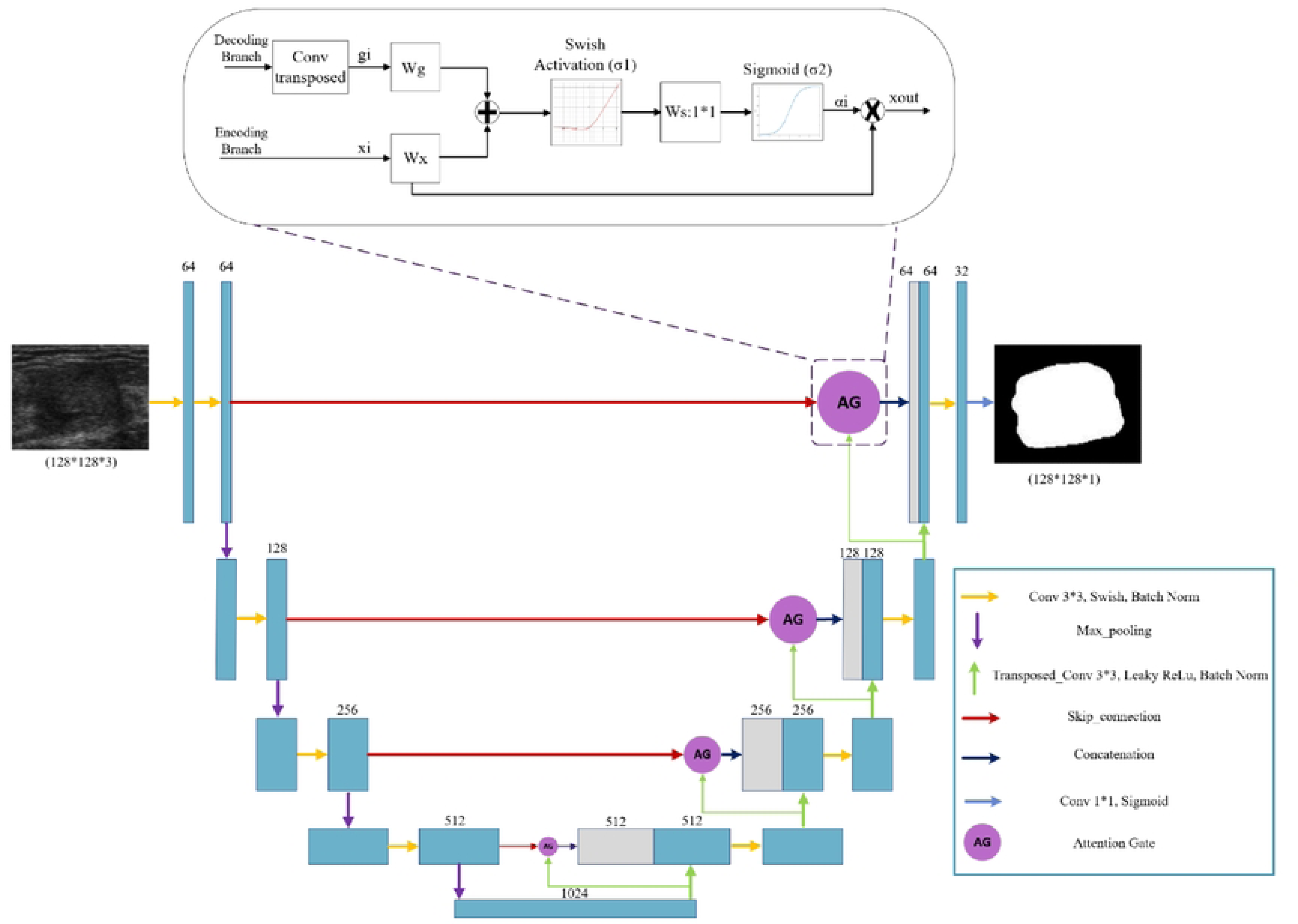
Bottom: The attention UNet architecture inspired by [49] the AG selectively emphasizes important features and suppresses irrelevant features. Top: Illustration of the proposed additive Attention Gate (AG), which employs a gating signal (g) derived from applying transposed convolution on coarser scale features and the features from the encoding path to analyze the activations and contextual information for selecting spatial regions.

**Figure 4.**
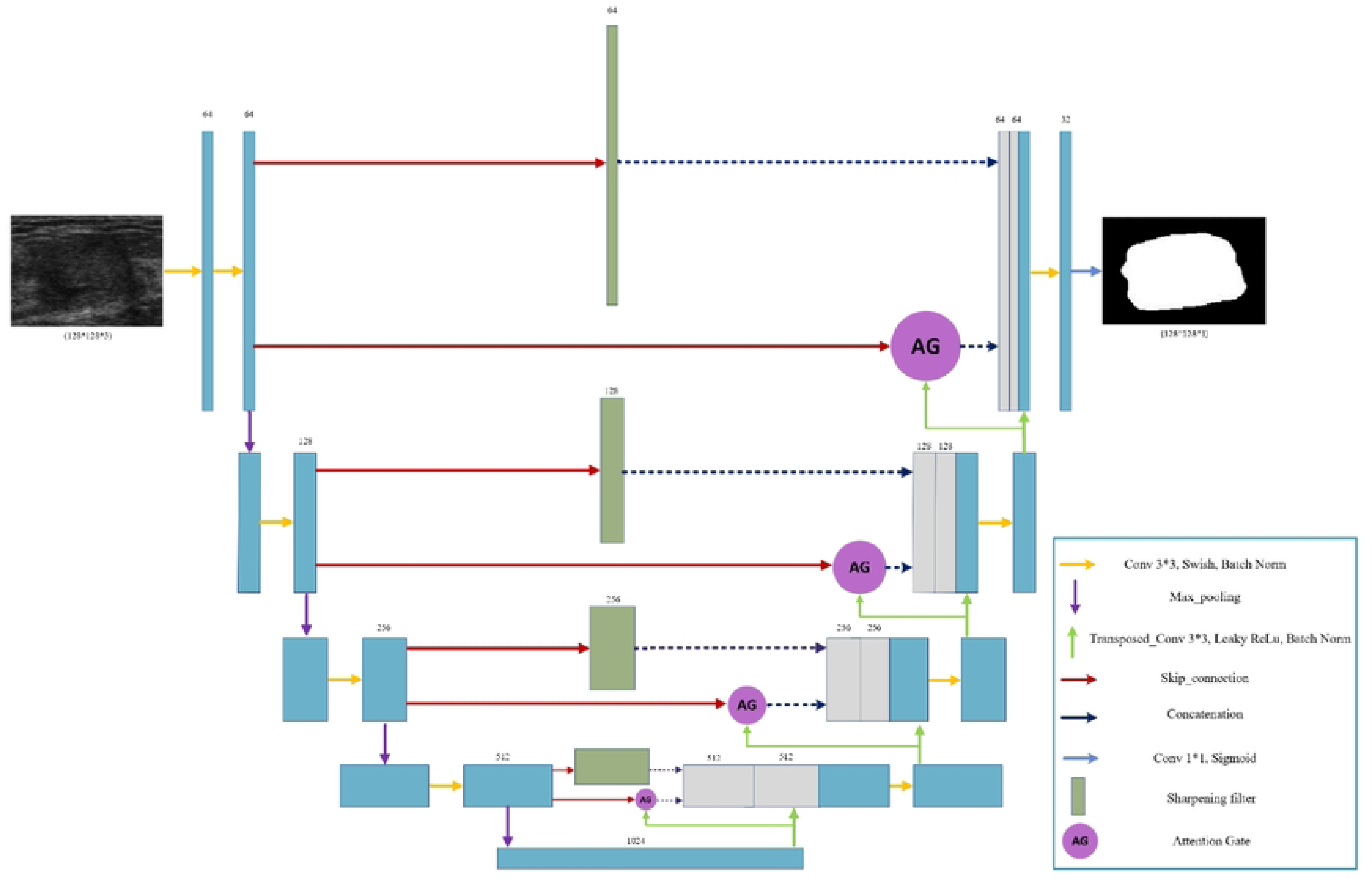
the proposed Sharp Attention UNet architecture. The combination of features enhances the network’s ability to capture details and context, leading to improved feature representation and segmentation performance.

The UNet architecture, proposed by Ronneberger et al.[44], resolves the issue of decreased fine-grained spatial details in the decoder section by incorporating skip connections that directly link corresponding layers of the encoder and decoder [45]. The UNet architecture is a powerful and flexible model for medical image segmentation. Its ability to handle limited training data, accurately identify regions of interest, handle variations and noise, and preserve spatial information makes it highly suitable for a wide range of medical imaging applications[46]. The UNet architecture is based on the fully convolutional

network approach, which enables end-to-end pixel-wise predictions. It consists of two main parts: the contracting path (encoder) and the expansive path (decoder). These two parts are connected to each other by skip connections. These skip connections help in preserving the spatial details that are usually lost during the downsampling process in the encoder. By combining the information from the encoder and decoder through concatenation, the spatial information is preserved while enhancing the depth of the feature maps. This enables the network to learn and understand the spatial relationships between different features more effectively.

#### 3.2.1 Sharp UNet

The simple skip connections used between the encoder and decoder within a UNet model can cause the gradients to vanish[47], which hampers the model’s capacity to accurately segment objects in images. Additionally, these simple skip connections may extract redundant low-level features and fail to capture multi-scale features or selectively focus on important regions in the input image. One approach to handle the loss of information due to the simple skip connections is applying sharpening filter on the encoder path features then concatenate the results with the decoder path [48]. The use of the sharpening filter layer allows for semantically fusing less dissimilar features.

In order to minimize the disparity between encoder and decoder features, in the Sharp UNet a sharpening filter applied within each pathway connecting the encoder and decoder. Image sharpening highlights intensity transitions in an image by using convolution with specific kernels or masks, such as the Laplacian filter. This filter captures changes in intensity across the image and enhances image details.

Figure 2 shows the sharp UNet architecture used in this paper. Among different Laplacian kernels, the kernel given in eq.10 presented the best performance in our application. Moreover, we observed an enhancement in the performance of our model by eliminating the final sharpening filter that was originally between the last encoder layer and the initial decoder layer.

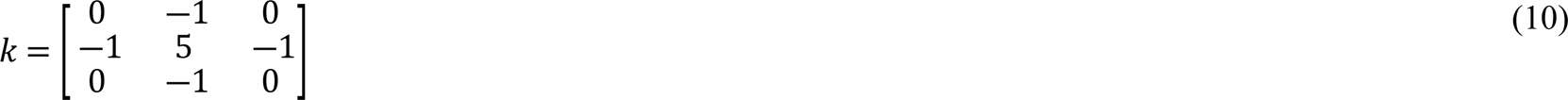

While sharp UNet shows a vast potential for image segmentation, it also comes with certain limitations. The risk of over-sharpening might lead to noise amplification or artifact introduction.

#### 3.2.2 Attention UNet

The attention mechanism enhances important features in neural network models. In[49], a new UNet algorithm with an attention module is proposed for pancreas segmentation in CT images. The network includes an encoder module for feature extraction, an attention module to capture contextual information, and a decoder module to restore the concatenated feature maps. Recent studies have shown that combining attention gates (AGs) in DL models can enhance network performance [50, 51] In our study, we present an AG architecture, as depicted in Figure 3, which draws inspiration from the attention gate used in a previous work [49].

The AG unit in Figure 3 acts as a transitional component in the model architecture. It receives the inputs of the decoder and encoder branches. In the encoder branch, convolutional layers employ a hierarchical approach to extract high-level image features by processing local information layer by layer. This process results in the spatial separation of pixels based on their semantics in a higher-dimensional space. By sequentially processing local features, the model can incorporate information from a larger receptive field into its predictions. The feature-map 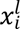 represents the extracted high-level features from layer *l* with the subscript *i* denoting the spatial dimension for each pixel’s value (*a*_*i*_, *x*_*i*_, and *g*_*i*_ in Figure 3).

The AG mechanism can be expressed in eq.11, and 12.

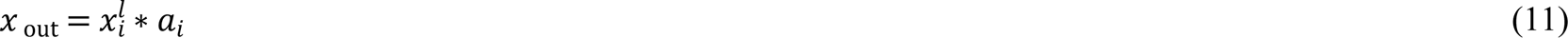

where *x*_out_ is the feature map calculated by element-wise multiplication between the input feature map inline1 and attention coefficients *a*_*i*_, which *a*_*i*_ is expressed mathematically as:

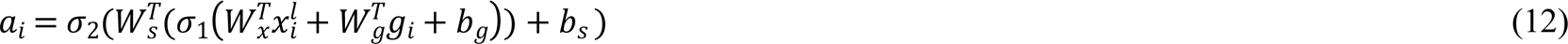

where σ_1_ is the swish activation function and σ_2_ is the sigmoid function. The feature map 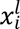 and the gating signal vector *g*_*i*_ undergo linear transformations using 1*1 channel-wise convolutions. The parameters *W*_*x*_, *W*_*g*_, and *W*_*s*_ are trainable, and the bias terms *b*_*g*_ and *b*_*s*_ are set to zero for simplicity. Experimental results suggest that setting these bias values to zero does not adversely affect the model’s performance.

#### 3.2.3 The Proposed Sharp Attention UNet

In this section, we develop a novel architecture by combining the Sharp UNet and Attention UNet models introduced earlier. The motivation behind this fusion is to utilize the advantages of both architectures and extract features that benefit from both the sharpening technique and the attention mechanism. The attention gate features, obtained from the Attention UNet, and the sharpened features derived from the Sharp UNet, are concatenated with the decoder features. This approach aims to enhance the network’s capability to capture fine details and relevant contextual information, thus improving the overall feature representation and segmentation performance. The proposed architecture is depicted in Figure 4. In this architecture, we employ the identical sharpening filter and attention gate (AG) mechanism as those introduced in the previously described Sharp UNet and Attention UNet architectures.

## 4. Results

Within this section, we assessed the efficiency of the proposed models through the application of a range of performance metrics, including the Dice coefficient, accuracy, loss, Dice loss, precision, sensitivity, specificity, F1, and recall. These metrics are explained in detail in appendix A. The evaluation was carried out on the BUSI dataset. The proposed model was developed using Python 3.7 with Keres API and was trained and tested on NVIDIA RTX A3000 GPU, and NVIDIA Tesla P100 GPU. The model underwent training for 300 epochs using the Adam optimizer (learning rate = 0.001, beta1= 0.9, beta2= 0.9, epsilon=0.0000001).

We randomly divided the BUSI dataset into training, validation, and test sets, with proportions of 80% (624 images), 10% (78 images), and 10% (78 images) respectively. The training set is used to optimize the model’s parameters by adjusting them based on input data and corresponding target values. After each epoch, the model’s performance is evaluated on the validation set to fine-tune hyperparameters such as learning rate, layer count, and neuron count per layer. This process, known as hyperparameter tuning, aimed to optimize performance. The best model is decided based on the validation results and saved as the optimal model. To evaluate performance, a separate test set that was not part of the training or validation processes is utilized.

For the analysis we tested the performance of our proposed Sharp Attention UNet with different input resolutions, as well as an analysis of various activation functions. Furthermore, the impact of applying CLAHE as a preprocessing step is explored. In addition, the proposed model’s performance is compared to other models presented in this paper which were evaluated using the same dataset.

### 4.1 Quantitative analysis of the proposed Sharp Attention UNet model for different input image resolutions

Segmentation was performed using different network input sizes (32*32, 64*64, and 128*128) for our proposed Sharp Attention UNet. However, due to excessive memory usage by the GPU, attempts to increase image resolutions with a batch size of 32 were not possible. The experiment was performed for 300 training epochs with a batch size of 32. CLAHE is applied as a preprocessing step. The results of the experiment, presented in Table 1, include performance metrics calculated using equations in appendix A. Analysis of the performance metrics in Table 1 reveals that the input size of 128×128 achieves the highest values.

**Table 1.** Quantitative Evaluation of Sharp Attention UNet of different input sizes on the BUSI dataset.

| Input resolution | Batch_size | Accuracy | Loss | Dice loss | Precision | F1 | Sensitivity | Specificity | Dice coefficient |
| --- | --- | --- | --- | --- | --- | --- | --- | --- | --- |
| 32*32 | 32 | 0.9737 | 0.2738 | 0.1381 | 0.9076 | 0.8683 | 0.8343 | 0.9913 | 0.8619 |
| 64*64 | 32 | 0.9804 | 0.1879 | 0.0962 | 0.9231 | 0.9097 | 0.8996 | 0.9942 | 0.9038 |
| 128*128 | 32 | **0.9792** | **0.1086** | **0.0717** | **0.9412** | **0.9412** | **0.9426** | **0.9936** | **0.9283** |

### 4.2 Quantitative analysis of the proposed Sharp Attention UNet model for different activation functions

The quantitative analysis conducted on the Sharp Attention UNet model with various activation functions (ReLu, LRelu, swish, and mish) is presented in Table 2. The results indicate that swish outperforms the other activation functions. This suggests that the swish activation function is more suitable for the Sharp Attention UNet model compared to the alternative activation functions that were evaluated.

**Table 2.** Quantitative evaluation of the Sharp Attention UNet with different activation functions.

| Activation Function | Accuracy | Loss | Dice loss | Precision | F1 | Sensitivity | Specificity | Dice coefficient |
| --- | --- | --- | --- | --- | --- | --- | --- | --- |
| ReLU | 0.9797 | 0.2079 | 0.1043 | 0.9158 | 0.9006 | 0.8885 | 0.9912 | 0.8957 |
| LRelu(alpha=0.1) | 0.9809 | 0.1738 | 0.0957 | 0.9330 | 0.9101 | 0.8906 | 0.9937 | 0.9043 |
| Swish | **0.9792** | **0.1086** | **0.0717** | **0.9412** | **0.9412** | **0.9426** | **0.9936** | **0.9283** |
| Mish | 0.9724 | 0.1234 | 0.0906 | 0.9121 | 0.9204 | 0.9126 | 0.9822 | 0.9104 |

### 4.3 Quantitative analysis of the proposed Sharp Attention UNet model before and after CLAHE preprocessing

Figure 5 presents a visual representation of two breast images before (a) and after (b) CLAHE enhancement. The samples depicted in the figure comprise two samples from the BUSI dataset. By comparing these images, we can observe the extent to which the CLAHE method enhances the contrast of the images from both datasets. In the results section we will show the performance of the segmentation model with and without CLAHE contrast enhancement.

**Figure 5.**
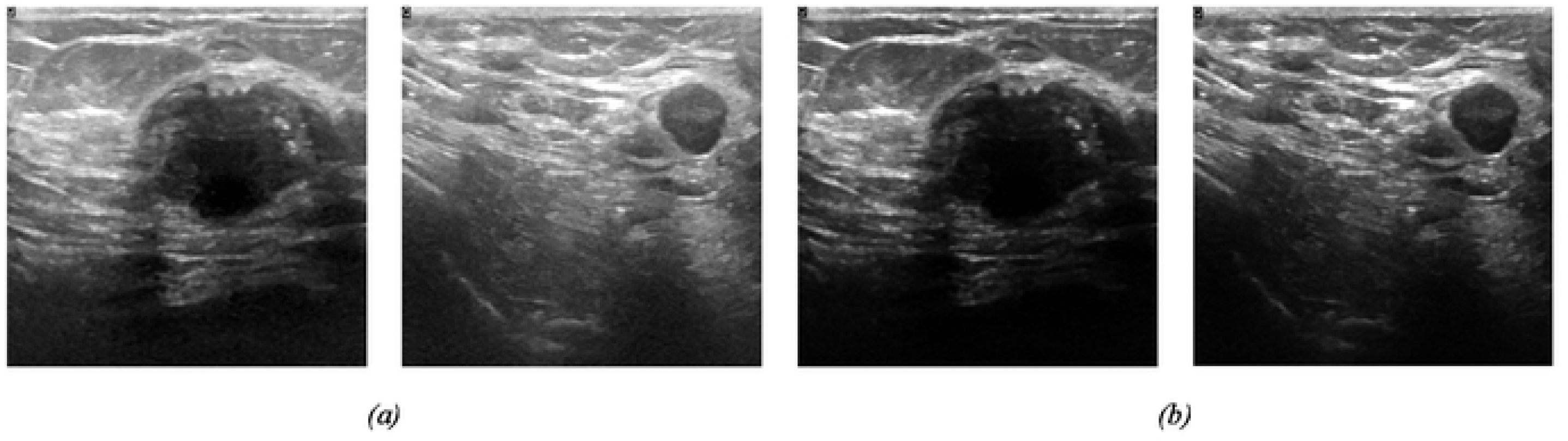
(a) Original image (b) Adaptive contrast image from CLAHE

To assess the effectiveness of CLAHE as the preprocessing technique, the proposed Sharp Attention UNet model was evaluated on the test set using an image size of 128*128 and a batch size of 32. The swish activation function was employed as the activation function. The evaluation aimed to determine the model’s efficacy by assessing both the original and CLAHE-enhanced versions of the BUSI dataset. The findings presented in Table 3 demonstrate that the application of CLAHE has a positive effect on the results.

**Table 3.** Quantitative evaluation of applying CLAHE TO the Sharp Attention UNet.

| Dataset | Accuracy | Loss | Dice loss | Precision | F1 | Sensitivity | Specificity | Dice coefficient |
| --- | --- | --- | --- | --- | --- | --- | --- | --- |
| Original BUSI | 0.9801 | 0.2126 | 0.1067 | 0.9086 | 0.8990 | 0.8923 | 0.9913 | 0.8940 |
| Enhanced BUSI | **0.9792** | **0.1086** | **0.0717** | **0.9412** | **0.9412** | **0.9426** | **0.9936** | **0.9283** |

### 4.4 Comparative Analysis of Neural Network Models: Performance Evaluation and Advantages of the Proposed Architecture

Tables 4 displays the performance metrics achieved by four distinct neural network models presented in this study. These models are built upon previous works, namely UNet[44], Attention UNet[49], and Sharp UNet [48]. While the primary network architectures for the first three models have been previously introduced in existing literature, the same backbone architecture and hyperparameters were used for all four models which the detail were discussed in detail in previous sections in this paper. The image input size of 128*128 was standardized, along with the application of augmentation techniques and CLAHE preprocessing. The optimization technique, batch size, loss function, and learning rate were kept consistent for all models. The results of these comparisons revealed that the proposed architecture outperformed the other models across all validation parameters.

**Table 4.** Comparison of the performance of the Sharp Attention UNet model with other UNet based models on the BUSI dataset.

| Model | Accuracy | Loss | Dice loss | Precision | F1 | Sensitivity | Specificity | Dice coefficient |
| --- | --- | --- | --- | --- | --- | --- | --- | --- |
| UNet | 0.9817 | 0.2311 | 0.1390 | 0.9115 | 0.8672 | 0.8336 | 0.9937 | 0.8610 |
| Attention UNet | 0.9837 | 0.1662 | 0.1061 | 0.9241 | 0.8999 | 0.8780 | 0.9936 | 0.8939 |
| Sharp UNet | **0.9838** | 0.1861 | 0.1234 | 0.8846 | 0.8856 | 0.8904 | 0.9919 | 0.8766 |
| **Proposed model** | 0.9792 | **0.1086** | **0.0717** | **0.9412** | **0.9412** | **0.9426** | **0.9936** | **0.9283** |

The outcomes achieved by the suggested algorithm indicate enhanced sensitivity on the dataset, with a notable improvement of more than 5% compared to the second-ranked model. Moreover, the Dice coefficient also experienced a noticeable increase of almost 3% compared to the second-best model.

Through experimental evaluations, we demonstrate in Tables 4 that our model outperforms existing state-of-the-art methods in accuracy, robustness, and computational efficiency.

We applied McNemar’s statistical test to compare the dichotomous performance of each pairwise segmentation model evaluated in this paper. McNemar’s test can evaluate the statistical significance of differences between the segmentation models[52, 53] under reasonable assumptions. The test was performed on a pixel-wise basis for the entire test set comprising 78 images with dimensions of 128*128 pixels. For all the images in the test set, the number of discordant entries was calculated by comparing the segmentation results of each two models while keeping the ground truth mask as the true value. The obtained McNemar’s test values are compared to the chi-squared distribution with 1 degree of freedom. The resulting p-value was calculated to determine the statistical significance of the observed differences between the models. The commonly used alpha value is considered 0.05 (5%), indicating that a p-value below 0.05 was considered significant. However, as our database is small, we considered 90% confidence level (α=0.1). However, when conducting multiple statistical tests simultaneously, such as comparing multiple pairs of models, there is an increased chance of obtaining false positives. To address this, the Bonferroni correction adjusts the alpha value to maintain an appropriate level of significance [54].[55]. As shown in Table 5, the comparison between the Attention UNet and Sharp UNet models yielded the highest p-value, suggesting a lack of statistically significant distinction between these two models in terms of performance. Our proposed model, which incorporates both Attention gates and sharpening filters within a UNet framework, demonstrated a significant improvement when compared to the earlier models. However, with the considered alpha value in this study, there is not a significant difference between the proposed model and attention UNet designed in this work. Still, on the parametric evaluation tests, the proposed model shows better results. This finding underscores the synergistic effects of integrating attention mechanisms and sharpening filters, resulting in enhanced performance, and indicating the added value of our proposed model over the standalone Attention UNet and Sharp UNet models.

**Table 5.** p values of the McNemar’s test for comparing model evaluation. Values under 0.02 imply the error distribution from the two compared models are significantly different after Bonferonni correction.

| Model | UNet | Attention UNet | Sharp UNet |
| --- | --- | --- | --- |
| UNet |  |  |  |
| Attention UNet | <0.02 |  |  |
| Sharp UNet | 0.045 | 0.079 |  |
| Proposed model | <0.02 | 0.023 | <0.02 |

## 5. Discussion

In this paper we fine-tuned and evaluated the UNet, Sharp UNet, and Attention UNet models for breast ultrasound segmentation. The reason why we choose these networks is the potential of UNet-based models for segmenting medical images. These models, which are already established in the literature, were further optimized, and fine-tuned using various techniques to enhance their performance metrics. The application of optimization techniques plays a crucial role in improving the segmentation accuracy of the CNN-based models. By carefully selecting and fine-tuning these techniques, we observed notable improvements in the performance metrics.

The results highlight the significance of optimization techniques and fine-tuning in improving the efficiency of UNet-based models for breast ultrasound segmentation. By carefully considering and optimizing various aspects of the models, we were able to achieve improvements in performance metrics, demonstrating the potential of these models for accurate and reliable segmentation of breast ultrasound images.

Moreover, the study explored the combination of Sharp UNet and Attention UNet to create a novel model, known as Sharp Attention UNet. This hybrid architecture demonstrated superior performance compared to individual models, emphasizing the potential benefits of integrating different network components to increase their respective performance. The advantages of the introduced model over the previous similar architectures is its ability to extract more meaningful features by incorporating sharpening filter and attention gate module. Sharp Attention UNet has several possible uses in clinical practice. It could both potentially replace an experienced radiologist when one is not available or assist a radiologist to improve diagnostic accuracy. There is also a powerful potential to combine Sharp Attention UNet with standardized image acquisition to allow rapid, automatic diagnosis without a radiologist or sonographer. One such approach incorporates the use of AI with volume sweep imaging (VSI). VSI is an imaging technique in which an individual without prior ultrasound training performs blind sweeps of the ultrasound probe over a target region such as the breast. VSI has already been clinically tested for breast, lung, thyroid, right upper quadrant, and obstetrics indications [6, 9, 56–62]. Individuals have been shown to be able to perform VSI after only a few hours of training [63, 64]. Integration of VSI with artificial intelligence has already shown promising results for breast and obstetrics indications [12, 65]. Future study testing the performance of Sharp Attention and VSI would be a promising step toward developing rapid and automatic diagnosis of breast lumps without a radiologist or a sonographer.

While the results are promising, it is important to acknowledge the limitations of this study. The evaluation was performed on a specific dataset with a limited number of images, and there may be variations in image quality and characteristics across different datasets. Therefore, further investigation on larger and more diverse datasets is necessary to assess the generalizability and robustness of the proposed models. Furthermore, the optimization techniques utilized in this work should be studied on other segmentation models to find their limitations and finding replacement solutions.

In future research, it would be also beneficial to explore additional optimization techniques, such as different data augmentation strategies or advanced regularization methods, on different models to further enhance the performance of the segmentation models. Additionally, investigating the transferability of these models to other medical imaging tasks or modalities could expand their potential applications and impact.

## 6. Conclusion

This paper presents a comparative study that explores the impact of different factors such as image preprocessing and various optimization techniques on the performance of UNet, Sharp UNet, and Attention UNet models for breast ultrasound image segmentation. We also introduced a novel UNet-based model, Sharp Attention UNet, by combining Sharp UNet and Attention UNet models. Our results show the potential of Sharp Attention UNet. We showed a dice coefficient, specificity, sensitivity, and F1 score of 0.92, 0.99, 0.94, and 0.94 respectively. Furthermore, McNemar’s test results show significant improvement when we compared our model against UNet, and Sharp UNet. This could be used to develop rapid automatic diagnosis of breast lumps without a radiologist or sonographer. Since most of the people in the world lack access to any form of medical imaging this potentially lifesaving artificial intelligence could be a promising avenue to improving global health and the diagnosis of breast cancers.

## Acknowledgments

This work was supported by National Institutes of Health Grants R21EB025290, and R21AG070331.

We are grateful to Prof. Tim Baran for his suggestions on statistical comparisons.

## Appendix A

### Dice coefficient

Dice coefficient is a measure of the similarity between two sets. It is defined as eq.9.

### Accuracy

Accuracy is a measure of the overall performance of a binary classification model and is calculated as the ratio of correctly predicted pixels to the total number of pixels (eq.15).

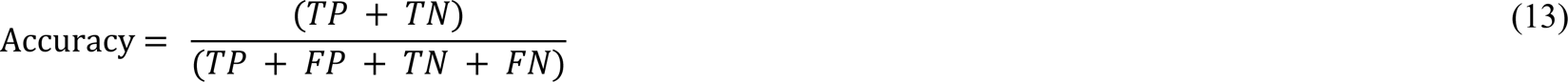

where True Positive (TP) is the number of pixels that are correctly classified as positive, True Negative (TN) is the number of samples that are correctly classified as negative, False Positive (FP) is the number of samples that are incorrectly classified as positive, and False Negative (FN) is the number of samples that are incorrectly classified as negative.

### Precision

Precision is a measure of the proportion of positive predictions that are correct and is calculated as the ratio of TP to the total number of predicted positives of each pixel.

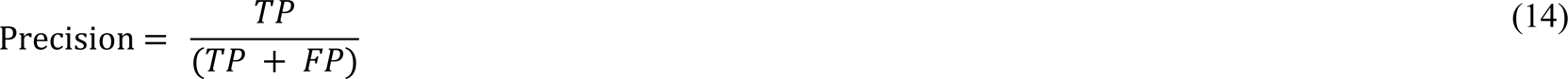

### Sensitivity

Sensitivity, also known as recall or True Positive Rate (TPR), is a measure of the proportion of actual positive pixels that are correctly identified as positive by the model and is calculated as the ratio of TP to the total number of actual positives of each pixel.

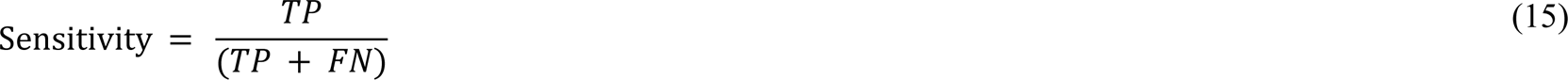

### Specificity

Specificity is a measure of the proportion of actual negative pixels that are correctly identified as negative by the model and is calculated as the ratio of TN to the total number of actual negatives of each pixel.

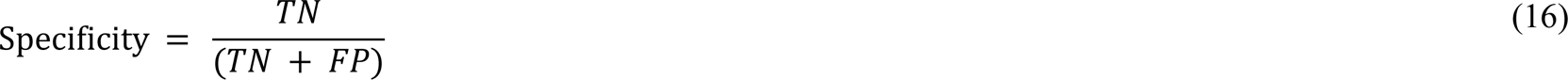

### F1 Score

The F1 score is a measure of the harmonic mean of precision and sensitivity and provides a balanced evaluation of the model performance.

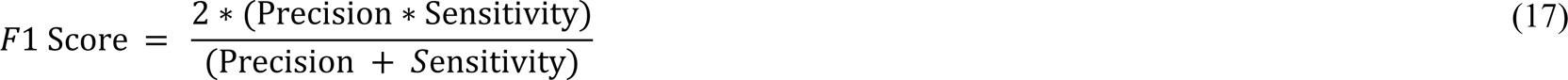

## Notes

### Competing Interest Statement

The authors have declared no competing interest.

